# Clustering single-cell RNA-seq data by rank constrained similarity learning

**DOI:** 10.1101/2021.04.12.439254

**Authors:** Qinglin Mei, Guojun Li, Zhengchang Su

## Abstract

**Motivation:** Recent breakthroughs of single-cell RNA sequencing (scRNA-seq) technologies offer an exciting opportunity to identify heterogeneous cell types in complex tissues. However, the unavoidable biological noise and technical artifacts in scRNA-seq data as well as the high dimensionality of expression vectors make the problem highly challenging. Consequently, although numerous tools have been developed, their accuracy remains to be improved.

**Results:** Here, we introduce a novel clustering algorithm and tool RCSL (Rank Constrained Similarity Learning) to accurately identify various cell types using scRNA-seq data from a complex tissue. RCSL considers both local similarity and global similarity among the cells to discern the subtle differences among cells of the same type as well as larger differences among cells of different types. RCSL uses Spearman’s rank correlations of a cell’s expression vector with those of other cells to measure its global similarity, and adaptively learns neighbour representation of a cell as its local similarity. The overall similarity of a cell to other cells is a linear combination of its global similarity and local similarity. RCSL automatically estimates the number of cell types defined in the similarity matrix, and identifies them by constructing a block-diagonal matrix, such that its distance to the similarity matrix is minimized. Each block-diagonal submatrix is a cell cluster/type, corresponding to a connected component in the cognate similarity graph. When tested on 16 benchmark scRNA-seq datasets in which the cell types are well-annotated, RCSL substantially outperformed six state-of-the-art methods in accuracy and robustness as measured by three metrics.

**Availability:** The RCSL algorithm is implemented in R and can be freely downloaded at https://github.com/QinglinMei/RCSL.

**Supplementary information:** Supplementary data are available at *Bioinformatics* online.

## 1 Introduction

Recent advances in single-cell RNA sequencing (scRNA-seq) technologies have revolutionized the study of many important biological processes, such as embryogenesis and tumorigenesis, in which an understanding of the functions and composition of heterogeneous cell types in the tissues is critical. As the transcriptome of a cell largely determines its molecular makeup, and thus its functions and cellular type, unsupervised clustering of individual cells based on their transcriptomes can be a powerful approach to identifying all the cell types including rare ones in complex tissues in an unbiased manner (Buettner, et al., 2015; Jiang, et al., 2016; Xu and Su, 2015). Despite great progress made in the last few years, the task remains highly challenging owing to the unavoidable biological noise and technical artifacts in scRNA-seq data as well as the high dimensionality of expression vectors.

The biological noise is related to the inherently stochastic nature of gene transcription in individual cells of the same type, due to the small copy number of molecules involved, unsynchronized cell cycles and uneven cell divisions (Becskei, et al., 2005; Kaern, et al., 2005; Paulsson, 2004; Raj and van Oudenaarden, 2008). As a result, even different cells of the same type display a broad range of variation in RNA levels (Bar-Even, et al., 2006; Newman, et al., 2006; Raj, et al., 2008; Taniguchi, et al., 2010; Xie, et al., 2008; Young, et al., 2012; Zenklusen, et al., 2008). Technical artifacts result from dropout events and batch factors in data generation, which are different from the conventional bulk RNA-seq data (Kiselev, et al., 2019). Intense efforts have been made to address these challenges over the past few years. For example, we previously proposed a quasi-clique-based algorithm with shared nearest neighbor(SNN), SNN-cliq, to identify groups of highly similar cells (Xu and Su, 2015). Guo *et al*. designed a top-to-toe pipeline (SINCERA) to distinguish major cell types (Guo, et al., 2015). Seurat, which has become one of the most popular choices for scRNA-seq data analysis, combines SNN graphs with Louvain community detection to group cells iteratively (Satija, et al., 2015). SC3 integrates the results of multiple clustering methods to obtain a consensus result (Kiselev, et al., 2017). Additionally, dimensionality reduction has also been integrated into clustering methods, such as pcaReduce (Žurauskienė and Yau, 2016) and ZIFA (Pierson and Yau, 2015), to reduce the computational complexity. Meanwhile, some approaches like CIDR, have been proposed to mitigate the impact of dropout events by improving the dissimilarity matrix (Lin, et al., 2017).

Another challenge in accurately clustering cells using scRNA-seq data is related to their high dimensionality. Although a cell may express tens of thousands of genes, only few of them determine its type (Graf and Enver, 2009), and we usually have no prior knowledge of which specific genes determine cell types. To address this, many new vector similarity metrics have been proposed such as SIMLR (Bo, et al., 2017) and MPSSC (Park, et al., 2018). SIMLR obtains a similarity matrix and identifies clusters via multikernel learning. MPSSC clusters a learned multiple doubly stochastic similarity matrix using sparse spectral clustering. In both the algorithms, a similarity matrix was learned from the data to better capture global structural relationships between cells. Nevertheless, these similarity metrics do not take into account local structures in quantifying cell similarities, which can be critical to discern subtle differences between cells of the same type and cells of different types, as indicated by our earlier proposed SNN metric (Xu and Su, 2015). In addition, most graph-based methods use the *k*-means algorithm to find clusters in a post-process step after constructing the similarity graph. Such two-stage algorithms inevitably lose some information from the original data in the post-process step.

In this work, we first propose a new metric that considers both global and local similarities between the expression vectors. Specifically, we use Spearman’s rank correlation to measure the global similarity and Neighbor Representation (NR) to capture the local similarity between the cells. NR has been successfully used in many dimensionality reduction algorithms, such as locally linear embedding (LLE) (Roweis and Saul, 2000), to capture local structures. We adaptively learn NR for each cell using an optimization procedure. Our metric is an optimized linear combination of Spearman’s correlation and the learned NR. Once the similarity matrix between cells is computed using this metric, the task of clustering the cells according to their types is to identify all block-diagonal submatrices in the similarity matrix or to partition the cognate similarity graph into several connected components, where each block-diagonal submatrix or connected component corresponds to a cell type. It has been proved that the number of block-diagonal submatrices in the similarity matrix is equal to the number of zero eigenvalues of its Laplacian matrix (Luxburg, 2007; Mohar, 1991; Nie, et al., 2016). Therefore, we posit that if we can reliably estimate the number of clusters or cell types *C* based on the similarity matrix, then we can transform it into a new matrix containing *C* block-diagonal submatrices, or partition the cognate similarity graph into *C* connected components, by constraining the rank of its Laplacian matrix. More specifically, if we can construct a matrix such that the number of zero eigenvalues of its Laplacian matrix is exactly equal to the estimated number of cell types, then we can divide cells into the same number of groups/types.

Based on this idea, we have developed a novel clustering algorithm RCSL (Rank Constrained Similarity Learning) by constructing a block-diagonal matrix such that the number of zero eigenvalues of its Laplacian matrix is equal to the estimated number of cell types in the dataset and its distance to the similarity matrix is minimized. When tested on 16 public scRNA-seq datasets with verified cell types, RCSL generally outperforms the six state-of-the-art methods compared.

## 2 Methods

### 2.1 The RCSL algorithm

RCSL takes a single-cell gene expression matrix ***M**_G′×N_* as the input, where *G’* denotes the number of genes, *N* denotes the number of cells, and an element *m_ij_* in ***M*** represents the expression value of gene *i* in cell *j*. RCSL consists of three steps (Fig. 1) detailed as follows.

**Fig. 1.**
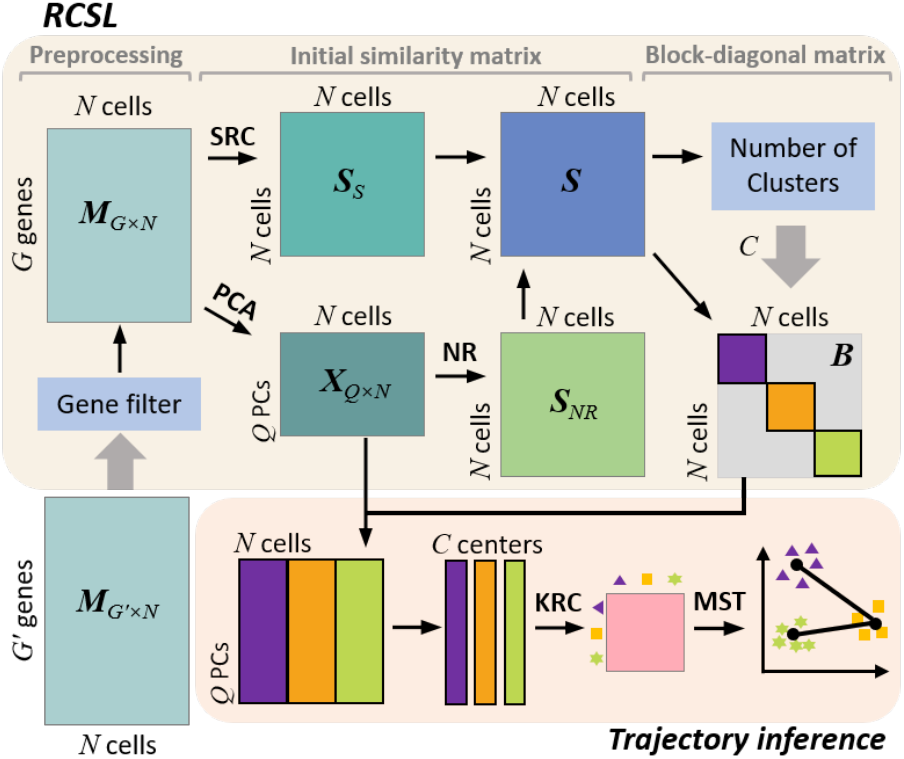
Overview of RCSL. First, given a gene expression matrix ***M**_G′×N_* as the input, RCSL filters out non-informative genes, resulting in matrix ***M**_G×N_*. Second, based on ***M**_G×N_*, RCSL computes Spearman’s rank correlation (SRC) matrix ***S**_S_* between the cells, performs PCA on the genes, and preserves the top *Q*-PCs as matrix ***X***. Third, RCSL learns the neighbour representation (NR) matrix ***S**_NR_* based on ***X***. Fourth, RCSL computes the similarity matrix ***S*** using a linear combination ***S**_S_* and ***S**_NR_*. Finally, RCSL estimates the number of clusters *C* in ***S***, and learns the block-diagonal matrix ***B*** from ***S,*** with the constraint that ***B*** has *C* zero eigenvalues. Each block-diagonal submatrix in ***B*** is a cluster of cells. RCSL also infers trajectories of the identified cell clusters/types.

#### Step 1. Data preprocessing

To control the quality of scRNA-seq data, we filter out rarely expressed genes and ubiquitously expressed genes, which contribute little to clustering. Specifically, we discard genes expressed in less than 2.5% of cells as well as genes expressed in > 97.5% of cells with a variance < 90% of the mean variance of these selected ubiquitously expressed genes. These parameters were chosen as we found that they could slightly improve the accuracy of clustering results on most of the datasets (Fig. S1). Let the resulting matrix be ***M**_G×N_*.

#### Step 2. Construction of similarity matrix

For each pair of cells *i* and *j* in ***M**_G×N_*, we calculate the Spearman’s rank correlation (SRC) between their expression vectors, defined as:

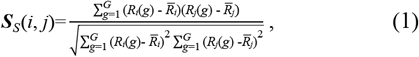

where *R_i_*(*g*) and *R_j_*(*g*) are the ranks of the expression level of gene *g* in cells *i* and *j*, respectively, and 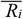 and 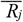 are the ranks in *i* and *j* of the gene in the middle ordered by gene IDs, respectively. Let the resulting matrix be ***S**_S_* (Fig. 1). As SRC is based on the ranks of values, it is insensitive to the difference of cell sizes. We perform principal component analysis (PCA) on ***M**_G×N_* (Wold, et al., 1987), and keep the top *Q* PCs that explain 95% of the variance. Let the resulting matrix be ***X**_Q×N_* (Fig. 1). We found that such dimensionality reduction has little influence on the accuracy of clustering (Fig. S1) but speeds up the algorithm somewhat (Fig. S2). Based on ***X**_Q×N_*, we compute the neighbour representation (NR) for each pair of cells as their local similarity as follows.

For each cell *i,* we find *k*-nearest-neighbours (*k*-NNs) using the Euclidean or cosine angle distance between their feature vectors in ***X**_Q×N_*. We use the Euclidean distance, as it generally performs better than the cosine angle distance in these study (see below). By default, *k* is set to 0.65*N*, which is a relatively large value to reduce the risk of information loss. Since cells of the same type are not necessarily close neighbours in Euclidean space as only few genes determine a cell’s type, in order to identify the close neighbours of the cells, we model each cell’s feature vector as a linear combination of the vectors of its neighbours; then those with higher weights are its closer neighbours. To find the weights, we solve a least-squares optimization problem:

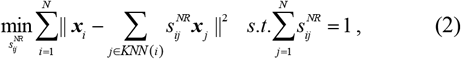

where ***x**_i_* is the feature vector of cell *i* in ***X**_Q×N_*, *KNN*(*i*) the *k-*NNs of *i* in Euclidean space, and 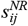 the weight of cell *j* on *i* 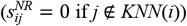.Intuitively, the greater the 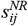 value, the more similar *j* is to *i*. Therefore, we call the learned weight vector 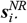 the NR of *i*. Let the resulting matrix for all the cells be 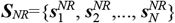. Then, we define the similarity matrix between the cells as a linear combination of ***S**_S_* and ***S**_NR_* (Fig. 1)

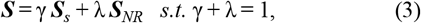

where γ ≥ 0 and λ ≥ 0 are scalar parameters that balance the contribution of ***S**_S_* and ***S**_NR_* in ***S***. By default, we set γ = 0.8, and λ = 0.2, as they generally perform best among other choices tested (see below).

#### Step 3. Calculation of block-diagonal matrix

We first estimate the number of clusters *C* by hierarchically clustering the cells based on ***S***, and find *C* that yields the largest Krzanowski-Lai index (Krzanowski and Lai, 1988) value from a range of *C* (by default, 4 to 12). However, for small datasets (N<3000), we use a two-step strategy to more accurately estimate *C.* Specifically, we choose three *C* values with the largest Krzanowski-Lai indexes, and pick the one with the largest sum of intra-class similarities based on ***S*** among the three clustering results of RCSL. The hierarchical clustering is performed using the R package *NbClust* with default settings.

To construct the block-diagonal similarity matrix ***B**_N×N_* between the cells, we adopt the Constrained Laplacian Rank (CLR) procedure (Nie, et al., 2010; Nie, et al., 2016). Briefly, CLR defines a diagonal matrix ***D**_B_* = diag (*d*_11_, *d*_22_,…, *d_NN_*), where 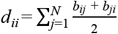 and *b_ij_* is the similarity between cell *i* and cell *j* in ***B***. The Laplacian matrix of ***B*** is defined as 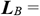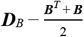. An important property of the Laplacian matrix is that the number of its zero eigenvalues equals the number of connected components in the graph defined by ***B*** (Fan, 1997; Mohar, 1991). Therefore, if a similarity matrix ***B*** can be found, such that the rank of its Laplacian matrix is exactly *N* − *C*, then ***B*** will have approximately *C* block-diagonal submatrix with proper permutations, and the corresponding similarity graph will contain *C* connected components. Each block-diagonal submatrix and corresponding connected component form a cluster of cells. Ideally, ***B*** should be highly similar to ***S***, and the rank of ***L**_B_* is exactly *N* − *C*. CLR therefore finds ***B*** by minimizing the difference between ***B*** and ***S,*** with the constraint that the rank of ***L**_B_* is *N* − *C*;

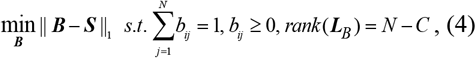

where the sum of each row in ***B*** is constrained to 1 to avoid rows of all zeros in ***B***. (see details in Supplementary Note).

### 2.2 Time complexity of the algorithm

Given the expression matrix ***M**_G′×N_*, with *N* cells and *G′* genes, Step 1 needs *N*\**G′* calculations, so it runs at O(*N*). In Step 2, we sequentially compute the SRC matrix ***S**_S_* and PCA matrix ***X**_Q×N_*, find *k*-NNs of *N* cells, and learn the ***S**_NR_* matrix, each of these procedures has a time complexity of O(*N*^2^). In Step 3, we perform *T* iterations (by default, *T* = 30) on the *N*Í*N* matrix ***S*** to estimate diagonal matrix ***B***, which needs O(*TN*^2^) calculations. As *T* is a small constant, Step 3 runs at O(*N*^2^). Therefore, the time complexity of the RSCL algorithm is O(*N*) + 5O(*N*^2^) = O(*N*^2^) (Fig. S3).

### 2.3 Inference of trajectory and pseudo-time

Based on the clustering results of RCSL, we infer the developmental trajectories and pseudo-temporal ordering of cells for time-series scRNA-seq data. For each identified cluster of cells, we compute its center as the mean of the feature vectors from one cluster in ***X**_Q×N_* (Fig.1), and the Kendall rank correlation (KRC) among all the centers. We construct a weighted similarity graph ***G***, where the vertices represent the centers and the edges represent their Kendall rank correlation values. We find the minimum spanning tree (MST) using the Prims’ algorithm (Prim, 1957). The MST that represents the shortest path connecting all the centers without any circles is the most parsimonious explanation of the relationships among the cell types during cell differentiation, and thus likely reflects their developmental trajectory. We determine the pseudo-temporal ordering between the cell types by using the distance from a cell type to the predefined starting cell type. The distance is defined as the reciprocal of the average similarity between the two types of cells in the similarity matrix ***S***.

### 2.4 scRNA-seq datasets

We collected 16 publicly available scRNA-seq datasets (Table 1), in which cell types were determined by the original authors using various methods, including microscopic inspections for the embryonic cells (oocyte, zygote, 2-cell stage, 4-cell stage, 8-cell stage,…, and blast cells), time of post-inductions for artificially induced differentiated cells (day 0, day 1, day 2,…), as well as molecular markers and cell purification using Fluorescence-Activated Cell Sorting (FACS) for the other datasets (for details see Table S1). To further ensure the accuracy of cell type annotations, we excluded cells with ambiguous labels including “dropped” cells in the Wang dataset, “contaminated” cells in the Xin dataset and “unclear” cells in the Muraro dataset. We normalize raw read counts of genes using CPM (counts per million) followed by adding a pseudocount of 1 and log (base 2) transformation.

**Table 1.**
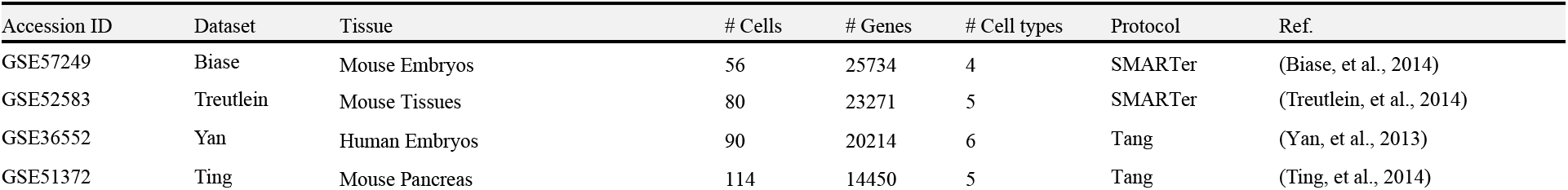
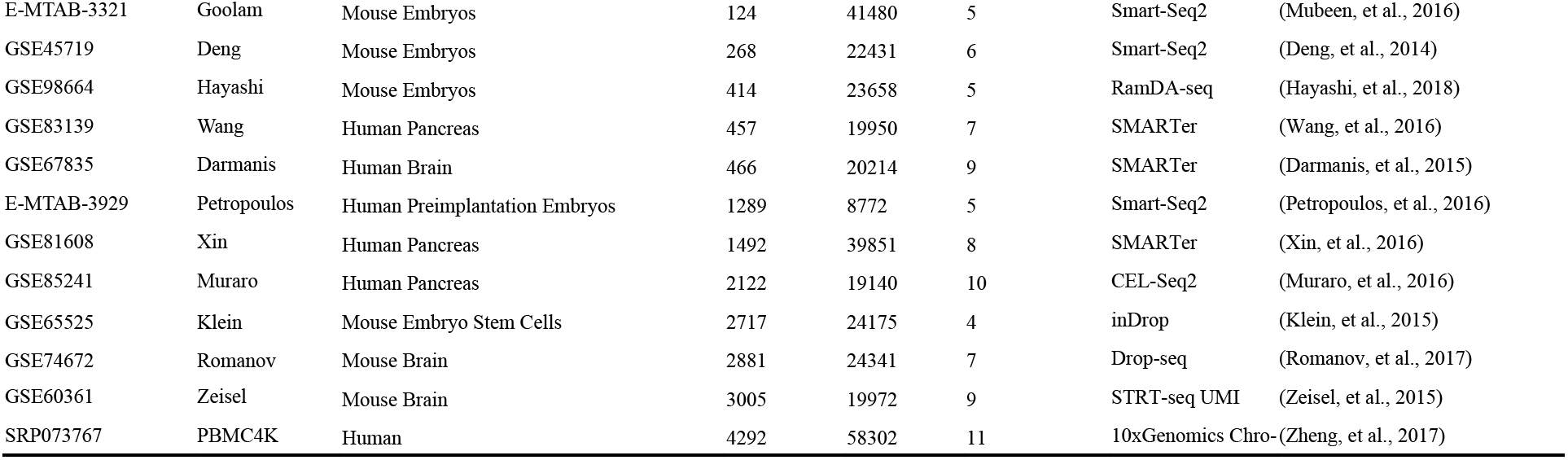
Summary of the 16 scRNA-seq datasets used in this study to assess the performance of the methods for clustering cells.

| Accession ID | Dataset | Tissue | # Cells | # Genes | # Cell types | Protocol | Ref. |
| --- | --- | --- | --- | --- | --- | --- | --- |
| GSE57249 | Biase | Mouse Embryos | 56 | 25734 | 4 | SMARTer | (Biase, et al., 2014) |
| GSE52583 | Treutlein | Mouse Tissues | 80 | 23271 | 5 | SMARTer | (Treutlein, et al., 2014) |
| GSE36552 | Yan | Human Embryos | 90 | 20214 | 6 | Tang | (Yan, et al., 2013) |
| GSE51372 | Ting | Mouse Pancreas | 114 | 14450 | 5 | Tang | (Ting, et al., 2014) |
| E-MTAB-3321 | Goolam | Mouse Embryos | 124 | 41480 | 5 | Smart-Seq2 | (Mubeen, et al., 2016) |
| GSE45719 | Deng | Mouse Embryos | 268 | 22431 | 6 | Smart-Seq2 | (Deng, et al., 2014) |
| GSE98664 | Hayashi | Mouse Embryos | 414 | 23658 | 5 | RamDA-seq | (Hayashi, et al., 2018) |
| GSE83139 | Wang | Human Pancreas | 457 | 19950 | 7 | SMARTer | (Wang, et al., 2016) |
| GSE67835 | Darmanis | Human Brain | 466 | 20214 | 9 | SMARTer | (Darmanis, et al., 2015) |
| E-MTAB-3929 | Petropoulos | Human Preimplantation Embryos | 1289 | 8772 | 5 | Smart-Seq2 | (Petropoulos, et al., 2016) |
| GSE81608 | Xin | Human Pancreas | 1492 | 39851 | 8 | SMARTer | (Xin, et al., 2016) |
| GSE85241 | Muraro | Human Pancreas | 2122 | 19140 | 10 | CEL-Seq2 | (Muraro, et al., 2016) |
| GSE65525 | Klein | Mouse Embryo Stem Cells | 2717 | 24175 | 4 | inDrop | (Klein, et al., 2015) |
| GSE74672 | Romanov | Mouse Brain | 2881 | 24341 | 7 | Drop-seq | (Romanov, et al., 2017) |
| GSE60361 | Zeisel | Mouse Brain | 3005 | 19972 | 9 | STRT-seq UMI | (Zeisel, et al., 2015) |
| SRP073767 | PBMC4K | Human | 4292 | 58302 | 11 | 10xGenomics Chro- | (Zheng, et al., 2017) |

### 2.5 Simulation datasets

We generated 10 simulated datasets containing 300~3,000 cells belonging to 4~7 cell types (Table S3) using the *Splatter* Bioconductor package (Zappia, et al., 2017). Each cell expresses 10,000-15,000 genes, whose levels are determined by the cell’s type. The script for constructing simulation datasets is available at https://github.com/QinglinMei/RCSL/tree/master/R.

### 2.6 Evaluation metrics

To measure the consistency between identified clusters and known cell types, we adopt three metrics: Adjusted Rand Index (ARI) (Hubert and Arabie, 1985), Normalized Mutual Information (NMI) (Strehl and Ghosh, 2002) and Fowlkes-Mallows index (FM) (Fowlkes and Mallows, 1983). We represent the known cell types as *R* and the identified clusters as *E*. Let *a* be the number of pairs of cells that are clustered in the same group in both *R* and *E*; *b* the number of pairs of cells that are clustered in the same group in *R* but in different groups in *E*; *c* the number of pairs of cells that are clustered in different groups in *R* but in the same group in *E*, and *d* the number of pairs of cells that are clustered in different groups both in both *R* and *E* (Kim, et al., 2019). Then, ARI, FM and NMI are defined as,

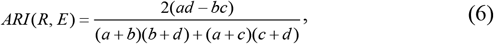

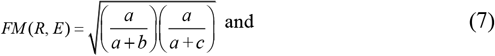

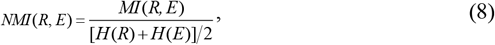

where *MI*(*R*, *E*) is the mutual information of *R* and *E*, and *H* an entropy function of *R* and *E*.

## 3 Results

### 3.1 Combination of global and local similarities improves the accuracy of RCSL

To find the optimal values of weights γ and λ of global similarity and local similarity, respectively, in the similarity metric ***S*** (formula (3)), and to see how they affect the accuracy of RCSL, we ran RCSL on the 16 datasets with varying vales of γ (0.0, 0.1,…, 1.0) and λ =1 − γ (1.0, 0.9,…, 0.0). As shown in Fig. 2, with the increase in γ, i.e., decrease in λ, the accuracy of RCSL generally increases, and reaches the highest level at γ = 0.8, λ = 0.2, then decreases. Thus, it appears that using only global similarity (γ = 1.0, λ = 0.0) as seen in most existing methods cannot guarantee the best performance on most datasets (Fig. 2). On the other hand, using local similarity alone (γ = 0.0, λ = 1.0) generally underperforms using global similarity alone (Fig. 2), due probably to information loss in the former. In this sense, it is not surprising that global similarity contributes more (80%) to the similarity score ***S*** than local similarity (20%) for the best performance. Nonetheless, this contribution of local similarity is necessary to the best accuracy in most datasets (Fig. 2). In addition, we evaluate the performance of Euclidean distance and cosine angle distance for defining *k*-NNs of the cells, and an approximate method LSH (Andoni, et al.) for finding the *k*-NNs, and find that Euclidean distance in combination with our method for finding *k*-NNs (Method) outperforms all other combination on most of the datasets (Fig. S4).

**Fig. 2.**
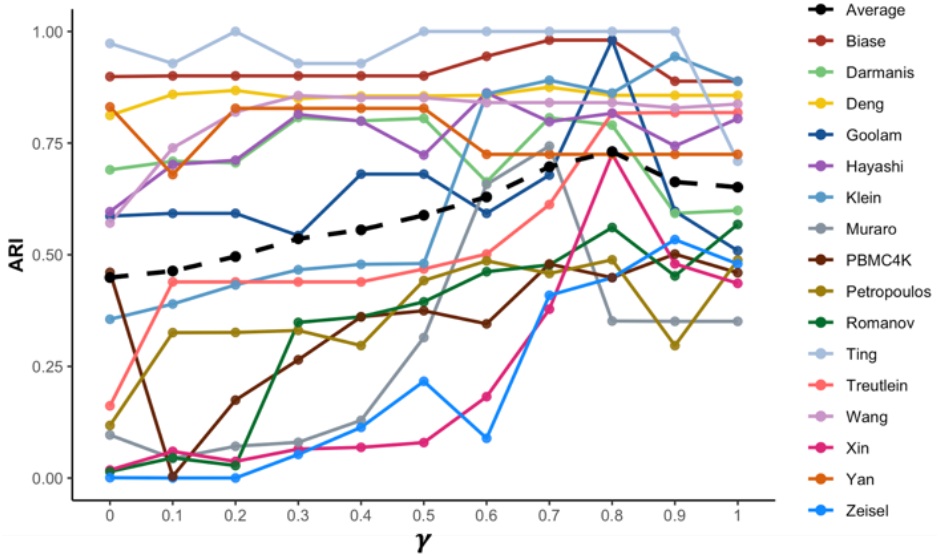
Effects of the values of γ and λ=1 − γ on the accuracy of RCSL.

### 3.2 RCSL outperforms existing methods in clustering cells

We next compare the performance RCSL on 16 scRNA-seq datasets with that of five state-of-the-art tools including SC3 (Kiselev, et al., 2017), SIMLR (Bo, et al., 2017), pcaReduce (Žurauskienė and Yau, 2016), CIDR (Lin, et al., 2017) and Seurat (Satija, et al., 2015) using three metrics (Methods). As mentioned before, SC3 is a popular method based on the consensus result of multiple methods; SIMLR is a similarity learning algorithm based on multikernel; pcaReduce is an agglomerative clustering method based on statistical modeling; CIDR is an ultrafast algorithm that imputes dropouts; and Seurat is widely applied to large datasets. Notably, both SIMLR and RCSL identify clusters by learning a block-diagonal similarity matrix from similarity matrices defined differently, therefore, we can compare their learned block-diagonal matrix. Moreover, to show the contribution of local similarity metric NR to the performance, we also implement a variant (RCSL2) of RCSL that does not use NR, thus is much faster than RCSL, though also runs at O(*N*^2^) (Fig. S3). More specifically, versions of the packages we compare with our algorithm are as follows: SC3 (package version 1.14.0 from Bioconductor); SIMLR (package version 1.12.0 from GitHub) (github.com/BatzoglouLabSU/SIMLR); pcaReduce (package version 1.0 from GitHub) (github.com/JustinaZ/pcaReduce); *k*-means (*kmeans* function built-in R version 3.6.0; CIDR (package version 0.1.5 from GitHub) (github.com/VCCRI/CIDR); and Seurat (package version 3.1.5 from CRAN). For PCA-Kmeans, we use our ***X**_Q×N_* as the input matrix. Since PCA-Kmeans is a stochastic algorithm, we run it 100 times and present the average of the results. In addition, since Kmeans cannot determine the number of clusters, we estimate the number of clusters by *NbClust* for Kmeans. For the other methods, we follow its corresponding instructions and tutorials provided by the authors and use its default parameters.

As shown in Fig. 3, both RCSL and RCSL2 outperforms the six existing methods on 11 of the 16 datasets (Table 1) based on ARI values (average 0.73 vs. 0.64). Specifically, on the Biase, Darmanis, Goolam, Treutlein Xin and Ting datasets, RCSL achieves significantly higher ARI than all the other algorithms. On the Deng and Romanov datasets, both RCSL and RCSL2 perform much better than the other algorithms. On the Wang and Hayashi datasets, RCSL also gains the highest ARI. Only on the Zeisel, Petropoulos, PBMC4K, Muraro and Yan datasets, SIMLR, Seurat or pcaRduce outperform RCSL (Fig. 3). However, on average, both RCSL and RCSL2 substantially outperform the six other methods (Fig. 3). Similar results are seen using NMI and FM (Figs. S5 and S6). These results indicate that both the optimized similarity metric (Fig. 2) and the clustering algorithm contribute to the outstanding performance of RCSL.

**Fig. 3.**
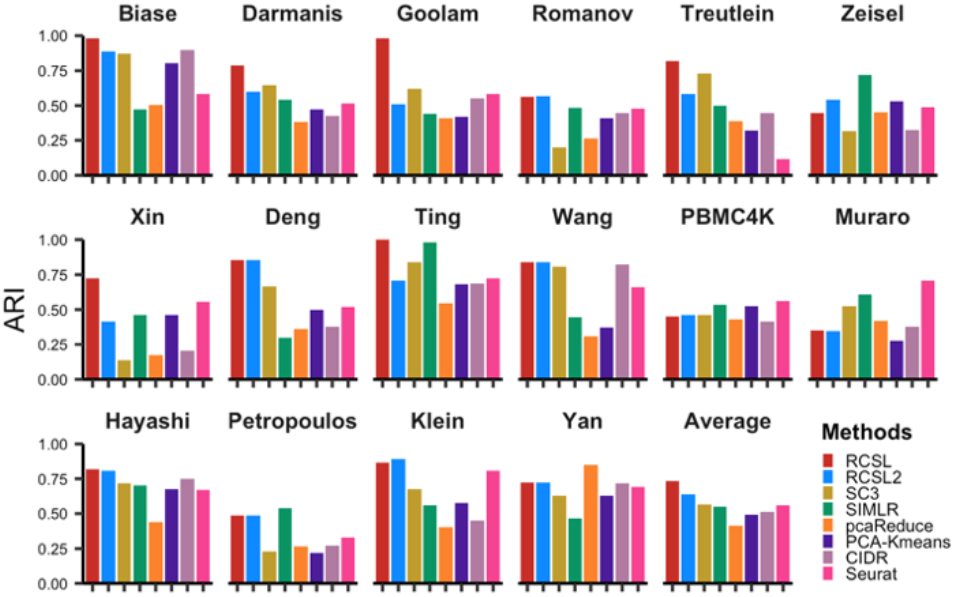
Performance of the algorithms (RCSL, RCSL2, SC3, SIMLR, pcaReduce, PCA-Kmeans, CIDR, Seurat) on the datasets measured by Adjusted Rand Index (ARI). The last panel shows the average ARI value for each algorithm over the 16 datasets

### 3.3 RCSL outperforms existing methods in learning the similarity among cells

We further seek to see whether the block-diagonal similarity matrices ***B*** learned by RCSL have the intended block-diagonal structures. To this end, we compare the SRC matrix ***S**_S_*, the similarity matrix ***S*** in RCSL as well as block-diagonal matrices learned by RCSL, RCSL2 and SIMLR. In the ideal case, if all cells are correctly clustered, then when cells are sorted by their types, the resulting matrix should exhibit clear-cut block-diagonal submatrices, in which similarities between cells of different types are zero while those between cells of the same types are non-zero. Fig. 4 shows the matrices learned by the four methods on eight datasets, in which cells are ordered according to their annotated types, and the results of the other eight datasets are shown in Fig. S7. Clearly, for most datasets except for the Ting, Klein, Wang, Zeisel and Yan datasets, there are no obvious block-diagonal submatrices for cells in ***S**_S_* or ***S*** (Figs. 4 and S7). In contrast, cells in the similarity matrices ***B*** learned by RCSL, RCSL2 and SIMLR possess clear-cut block-diagonal structures in all the datasets (Figs. 4 and S7). However, there are subtle differences among the results of the three algorithms. For the Treutlein and Klein datasets, both RCSL and RCSL2 correctly cluster the cells according to their annotated types, whereas SIMLR incorrectly groups multiple annotated cell types into one cluster. For the Biase, Goolam and Darmanis datasets, RCSL and RCSL2 correctly clustered the cells according to their annotated types, while SIMLR divides one type into multiple clusters. For the Goolam, Ting and Xin datasets, RCSL2 divides one cell type into multiple clusters, while clusters identified by RCSL are in better agreement with their annotated types, indicating the importance of including NR in the similarity metric. In addition, clusters found by RCSL are cleaner than those identified by RCSL2 and SIMLR in the off-diagonal blocks. As expected, the quality of the learned block-diagonal similarity matrices is consistent with clustering results measured by ARI, NMI and FM (Figs. 3, S5 and S6). Taken together, these results demonstrate that RCSL can better learn the block-diagonal similarity matrix of different cell types than the two other methods.

**Fig. 4.**
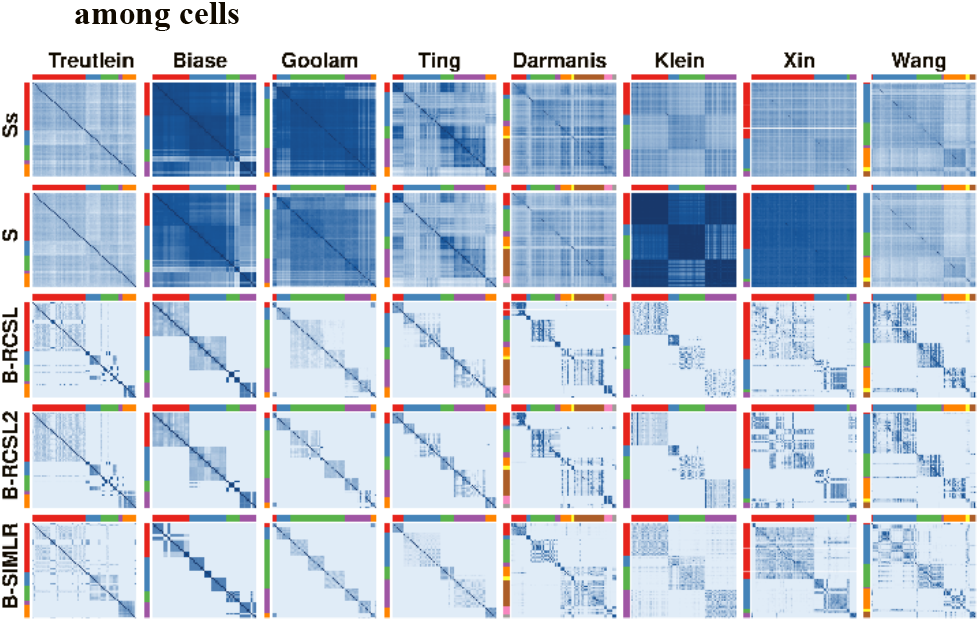
Heatmap of the Spearman’s rank correlation matrix ***S**_S_*, similarity matrix ***S*** in RCSL and the block-diagonal similarity matrices ***B*** learned by RCSL, RCSL2, SIMLR in the indicted eight datasets. Cells are arranged according to their annotated types indicated by the differently colored bar at the top and left of the matrices.

### 3.4 RCSL learns block-diagonal structures of cell-cell similarities in a step-wise manner

To see how RCSL gradually learns the block-diagonal structures of cell-cell similarities starting from a data matrix, thereby clustering cells, we visualize the data matrix ***M**_G′×N_*, global similarity matrix ***S**_S_*, similarity matrix ***S***, and block-diagonal matrix ***B*** of the 16 datasets using three visualization tools (PCA(Žurauskienė and Yau, 2016), *t*-SNE(Maaten and Hinton, 2008) and UMAP(McInnes, et al., 2018)). Fig. 5 shows the PCA 2D plots of the results from six datasets. The PCA plots for the remaining datasets as well as *t*-SNE and UMAP plots for all datasets are shown in Figs. S8, S9 and S10. Interestingly, in some datasets such as the Goolam, Ting, Klein and Yan datasets, most cells can be well separated according to their types simply by the data matrices ***M**_G′×N_* and ***S**_S_* in the 2D plots, owing to the markedly distinct gene expression patterns of different cell types. In other datasets such as the Treutlein and Muraro datasets, cells cannot be simply separated by the data matrices ***M**_G′×N_*, ***S**_S_* and ***S***. Particularly, for the Muraro and Klein datasets, cells of the same type are dispersed, while cells of different types are mixed in ***M**_G′×N_*, ***S**_S_* and ***S***. In contrast, different cell types in all the 16 datasets display clear-cut clusters in the block-diagonal matrix ***B*** (Figs. 5, S8, S9 and S10). Therefore, each step in the RCSL algorithm contributes to the identification of cell clusters/types in a dataset.

**Fig. 5.**
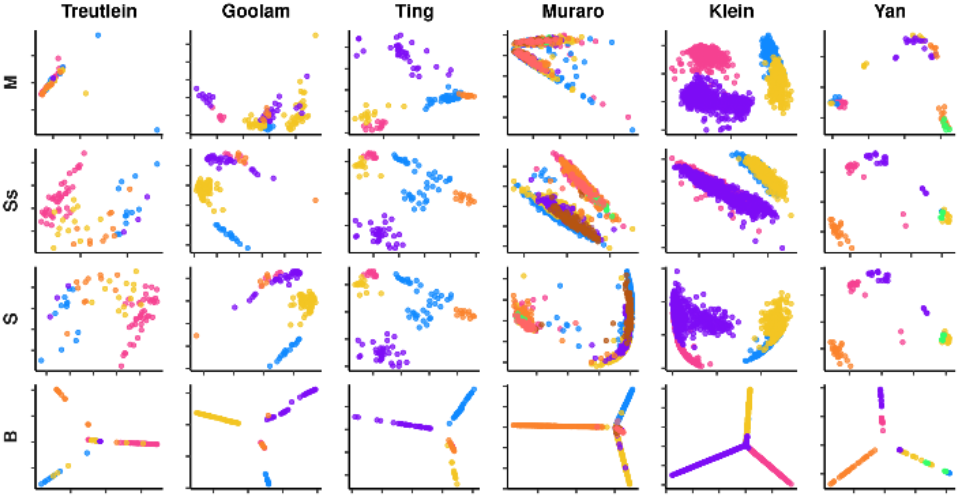
2-D PCA display of the expression data matrices and matrices produced by RCSL in the indicated datasets. The rows respectively correspond to ***M**_G′×N_*, Spearman’s correlation ***S**_S_*, similarity matrix ***S*** and block-diagonal matrix ***B***.

### 3.5 RCSL outperforms SIMLR in identifying sub-cell types

Notably, like SIMLR, RCSL also tend to divide one cell type into multiple sub-clusters in some datasets, particularly, developmentally related ones (Biase, Deng, Goolam, Klein, Hayashi, Petropoulos and Yan) (Figs. 4 and S7). This might reflect the hierarchical lineage relationships of cell types produced in cell differentiation processes. Three (Biase, Deng and Yan) of these datasets record subtypes produced during embryogenesis (Table S2). To see if RCSL is able to identify sub-cell types, we take a close look at the results from the three datasets (Fig. 6). For the Biase dataset containing four cell types, of which the Blast type is divided into the ICM and TE subtypes (Table S2), RCSL clusters the cells in five groups (Zygote, 2-cell, 4-cell, ICM and TE), and correctly splitting the Blast in the ICM and TE subtypes (Fig. 6). In contrast, SIMLR divides the cells into at least eight groups, with the 2-cell type incorrectly split in four clusters, the 4-cell type incorrectly split into three clusters (Fig. 6). Moreover, SIMLR fails to divide the Blast type into the ICM and TE subtypes (Fig. 6). For the Deng dataset containing six cell types, of which the Zygote type is further classified into Zygote and Early 2-cell types, the 2-cell type into Middle 2-cell and Late 2-cell types, and the Blast type into Early, Middle and Late types (Table S2), RCSL clusters the cells into five clusters, identifying the Middle and Late 2-cell types, but failing identify subtypes of the Zygote and the Blast type (Fig. 6). However, there might be no clear-cut difference among these subtypes based on their stages (Early, Mid and Late). In contrast, SIMLR divides the cells into at least 13 clusters, with the 8-cell type incorrectly split into four clusters, and the 16-cell type incorrectly split into at least two clusters, though it also correctly clusters the two subtypes of the 2-cell type (Fig. 6). For the Yan dataset containing six cell types, of which the Zygote type is divided into Oocyte and Zygote subtypes, RCSL clusters the cells in five groups, correctly identifying the 8-cell, 16-cell and Blast types, but failing to identify sub-types of the Zygote type. Although SIMLR also is able to correctly divide the Zygote type into two clusters, it splits the 16-cell type into two clusters, the Blast type into three clusters, and the 8-cell type into at least five clusters.

**Fig. 6.**
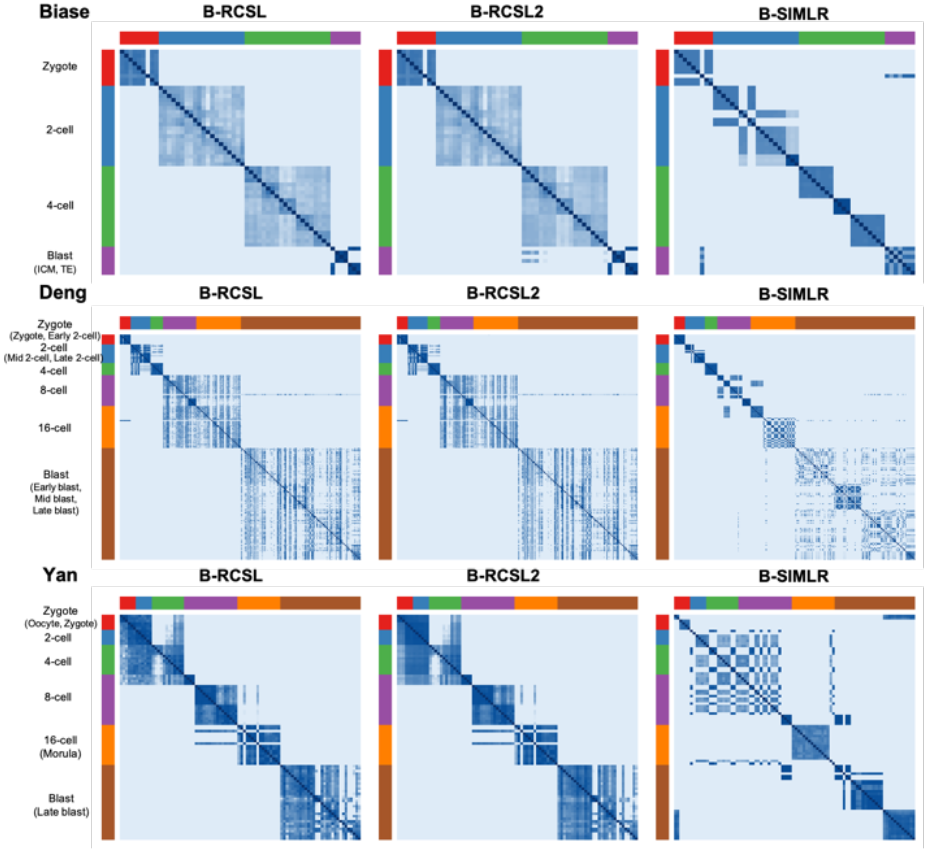
Heatmap of block-diagonal matrixes constructed by RCSL, RCSL2 and SIMLR for the Bias, Deng and Yang dataset where sub-cell types are recorded. Cells are arranged according to their annotated types and subtypes indicated by differently colored bars at the top and left of the matrices.

On the other hand, it is difficult to justify the subtypes identified by RCSL and SIMLR in the other datasets, as subtype information is unavailable. However, based on the results from the three datasets where some subtypes are classified, it appears that RCSL is more accurate in identifying sub-cell types than SIMLR that tends to over-cluster the cells.

### 3.3 RCSL achieves high clustering accuracy on simulated datasets

We have thus far demonstrated the high accuracy of RCSL for classifying cell types using the 16 datasets with well-annotated cell types. However, accurate cell type determination is still a challenging task, particularly, in larger datasets, it is difficult to guarantee 100% accuracy. To further evaluate the accuracy of RCSL, we ran it on 10 simulated datasets, in which the cell types are well defined (Table S3 and Method). Although RCSL slightly outperforms RCSL2 on the simulated datasets, both achieve a very higher average ARI of 0.95 and 0.94 (Fig. S11), respectively, suggesting that both are able to identify well-defined cell clusters. However, this almost same performance of RCSL and RCSL2 is in stark contrast to the result on the 16 real datasets, where RCSL substantially outperforms RCSL with an average ARI of 0.73 and 0.64, respectively (Fig. 3). Nevertheless, this is not surprising, as the clusters in the simulated datasets are clear-cut though the information is rather weak (Fig. S12), while the clusters in the real datasets are often vague with high background noise (Figs. 4 and S7). These results indicate that considering both local and global similarity is more critical to identify cell types when the data is highly noisy.

### 3.7 RCSL correctly infers trajectories and pseudo-time orders

Based on the clustering results, RCSL infers the developmental trajectories and pseudo-temporal orders of the identified cell types in a dataset, particularly when it is time-series-related. Fig.7 shows UMAP displays of the trajectories and pseudo-temporal orders of the identified cell types in four mouse embryo datasets (Goolam, Hayashi, Yan and Deng) and a human preimplantation embryos dataset (Petropoulos), in which both the cell types and developmental order are known. Remarkably, in each dataset, the inferred trajectory is the same as the real developmental trajectory, and the inferred pseudo-temporal ordering is also consistent with the true developmental stages. The results for the other datasets are shown in Fig. S13, where only an MST is shown if the starting cell type is unknown in a dataset, or it the dataset is not time series in nature.

**Fig. 7.**
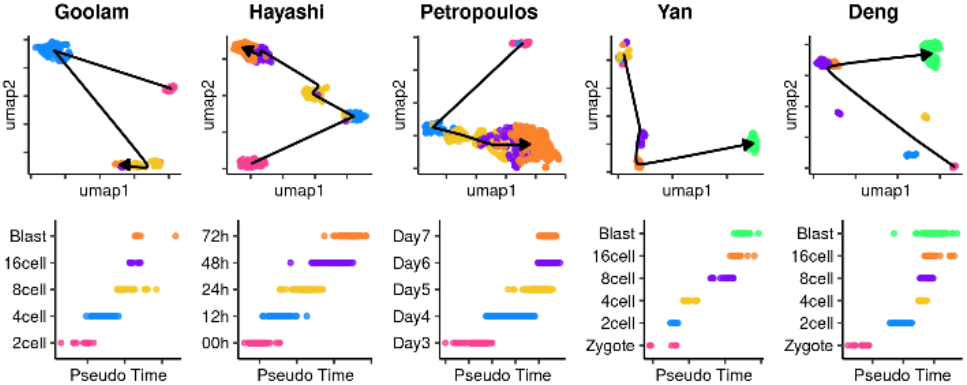
Inference of developmental trajectories and pseudo-temporal orders of the identified cell types in four mouse embryo datasets (Goolam, Deng, Yan, and Hayashi) and a human preimplantation embryos dataset (Petropoulos). The developmental trajectories (top) are visualized by UMAP. In the pseudo-temporal ordering of cells (bottom), the horizontal axis represents the estimated time of each cell type starting from the known initial cell type, and the vertical axis indicates the real cell stages/types.

## 4 Discussion

One of the major challenges for identifying heterogeneous cell types using scRNA-seq data lies in how to define the similarity among cells owing to inherent biological noise and unavoidable technical artifacts (Kiselev, et al., 2019). Moreover, only a few genes with similar expression patterns play a key role in defining a cell’s type (Graf and Enver, 2009). To tackle these problems, many similarity metrics have been developed (Kim, et al., 2018); however, they only consider global similarities (Bo, et al., 2017; Park, et al., 2018) even though local similarities can be crucial to differentiate the subtle difference between cells of the same type and cells of different types (Xu and Su, 2015). In this study, we propose a new metric that considers both the global similarity and local similarity between the cells. Specifically, we quantify a cell’s similarity to the other cells as an optimal linear combination of its global similarity and local similarity to other cells. For the global similarity of a cell, we utilize the SRC between the expression vector of the cell and those of all other cells in the data. For the local similarity of a cell, we adopt the NR that represents the cell’s feature vector (PC) as the optimal linear combination of the feature vectors of the cell’s *k*-NNs in the Euclidean or cosine angle distance space. Thus NR in effect adaptively adjusts the weights on the edges between the cell and its *k*-NNs in the corresponding similarity graph. The importance of incorporating NR into the similarity metric is demonstrated by the better performance of RCSL in almost all the datasets compared to RCSL2, which does not use NR, particularly, when the data are very noisy.

Another major challenge for identifying cell types using scRNA data is how to cluster cells by their types based on the similarity matrix(Kiselev, et al., 2019). Although many clustering algorithms have been proposed to identify cell types, the results are far from satisfactory due to the often complex structures of similarity matrices (Kiselev, et al., 2019). On the other hand, clustering cells in groups by their types is equivalent to converting the similarity matrix into a block-diagonal matrix by permutation and minimal adjustment of the similarity values. In the resulting block-diagonal matrix, each block-diagonal submatrix corresponds to a connected component in the corresponding similarity graph, i.e. a cluster or a cell type. We therefore adopt a method to compute such a block-diagonal matrix based on the similarity matrix. We first estimate the number of clusters defined in the similarity matrix and then iteratively find the block-diagonal matrix with the rank of its Laplacian matrix constrained.

We develop RCSL by combining the new similarity metric and the method for constructing the block-diagonal matrix, aiming to more accurately identify cells type in an often noisy scRNA-seq dataset. The results on the 16 diverse datasets show that RCSL substantially outperforms RCSL2 (a variant of RCSL that does not use local similarity), and RCSL2 outperforms the six other tools on many datasets, indicating that both the metric and block-diagonal matrix finding method contribute to the outstanding performance of RCSL. Although highly accurate, RCSL is limited by its O(*N*^2^) time complexity for computing ***S**_S_*, ***S**_NR_* and ***B***. We are currently developing a strategy to reduce the time complexity of RCSL to *N*log*N*, so that it can be applied to very large datasets generated from millions of cells.

## Acknowledgements

Q.M. conceived the algorithms and designed the software. Q.M, Z.S. and G.L wrote the paper. Z.S and G.L. oversaw the project. We would like to thank the three reviewers whose comments and suggestions improve the paper.

